# Single-cell tracing of the first hematopoietic stem cell generation in human embryos

**DOI:** 10.1101/750158

**Authors:** Yang Zeng, Jian He, Zhijie Bai, Zongcheng Li, Yandong Gong, Chen Liu, Yanli Ni, Junjie Du, Chunyu Ma, Lihong Bian, Yu Lan, Bing Liu

**Author notes:** These four authors contributed equally to this work (Yang Zeng, Jian He, Zhijie Bai, Zongcheng Li). Correspondence: Yu Lan and Bing Liu.

## Abstract

Tracing the emergence of the first hematopoietic stem cells (HSCs) in human embryos, particularly the scarce and transient precursors thereof, is so far challenging, largely due to technical limitations and material rarity. Here, using single-cell RNA sequencing, we constructed the first genome-scale gene expression landscape covering the entire course of endothelial-to-HSC transition during human embryogenesis. The transcriptomically defined HSC-primed hemogenic endothelial cells (ECs) were captured at Carnegie stage 12-14 in an unbiased way, showing an unambiguous arterial EC feature with the up-regulation of *RUNX1*, *MYB* and *ANGPT1*. Importantly, subcategorizing CD34^+^CD45^−^ ECs into CD44^+^ population strikingly enriched hemogenic ECs by over 10-fold. We further mapped the developmental path from arterial ECs via HSC-primed hemogenic ECs to hematopoietic stem progenitor cells, and revealed a distinct expression pattern of genes that were transiently over-represented upon the hemogenic fate choice of arterial ECs, including *EMCN*, *PROCR* and *RUNX1T1*. We also uncovered another temporally and molecularly distinct intra-embryonic hemogenic EC population, which was detected mainly at earlier CS 10 and lacked the arterial feature. Finally, we revealed the cellular components of the putative aortic niche and potential cellular interactions acting on the HSC-primed hemogenic ECs. The cellular and molecular programs and interactions that underlie the generation of the first HSCs from hemogenic ECs in human embryos, together with distinguishing HSC-primed hemogenic ECs from others, will shed light on the strategies for the production of clinically useful HSCs from pluripotent stem cells.

## Introduction

Hematopoietic stem cells (HSCs) give rise to all blood lineages and permanently maintain the adult hematopoietic system throughout the entire lifespan of an individual via self-renewal and differentiation. Although the ontogeny of HSCs during development has been extensively investigated in animal models including zebrafish and mouse, it is largely unclear in human embryos, given the limited accessibility of human embryonic tissues. By functional assessment in xenotransplantation, human HSCs are reported to be detected sequentially in multiple embryonic sites. Long-term multi-lineage repopulating HSCs are detected firstly in the aorta-gonad-mesonephros (AGM) region at Carnegie stage (CS) 14 (32 days post coitus, dpc), with a frequency of less than one per embryo equivalent, and then in the yolk sac several days later (CS 16, 35-38 dpc), showing an even lower frequency than that in AGM region^1, 2^. The first human HSCs manifest a phenotype of CD34^+^CD144^+^CD45^+^KIT^+^CD90^+^Endoglin^+^RUNX1^+^CD38^−/lo^CD45RA^−^, similar to those in the embryonic liver or cord blood^3–5^, although the enrichment is yet far from being efficient. An evident presence of HSCs in the embryonic liver, the major organ for HSC expansion during embryogenesis, is witnessed only after CS 17 (39-42 dpc), generally from 7-8 weeks of gestation^1, 2, 6^.

Histologically, the first HSCs are supposed to locate within the intra-aortic hematopoietic clusters (IAHCs) predominantly on the ventral wall of dorsal aorta (DA) in human embryos^3^. Given the expression of multiple endothelial markers in emerging HSCs, and the spatiotemporally intimate relationship between them and vascular endothelial cells (ECs), it has been widely proposed that human HSCs are derived from hemogenic ECs, a specified endothelial lineage with blood-forming potential, similar to what has been comprehensively validated in mice^7, 8^. Up-to-date, little is known about the developmental dynamics and molecular identity of the HSC-primed hemogenic ECs in human embryos, except for the simultaneous expression of endothelial feature genes and *RUNX1*, a transcription factor (TF) known to be expressed in hematopoietic-fated ECs and critically required for endothelial-to-HSC transition^9, 10^, and the lack of known hematopoietic markers such as CD43 (encoded by *SPN*) and CD45 (encoded by *PTPRC*).

Regarding the developmental path of hemogenic ECs that give rise to HSCs, most proposed views are established on the *in vitro* differentiation system of human pluripotent stem cells (hPSCs), which still fails to efficiently and reliably generate functional HSCs^11–14^. It is suggested that the hemogenic ECs and vascular ECs are specified early as distinct lineages from KDR^+^ mesoderm, and hemogenic ECs subsequently integrate with vascular ECs of dorsal aorta, undergoing endothelial-to-hematopoietic transition to form IAHCs^15, 16^. Most recently, activation of the arterial program in hemogenic ECs, which is regulated by MAPK/ERK and Notch signaling, has been reported to be required for the lymphopoiesis in PSC differentiation system, whereas the hemogenic ECs without arterialization demonstrate only primitive and myeloid-restricted hematopoietic potential^14, 17^. It remains elusive whether HSC-primed hemogenic ECs are derived from arterial ECs and what are the differences between the putative distinct hemogenic EC populations, if they exist, during human embryogenesis.

HSC development in AGM region needs the signals from surrounding niche cells, including at least sub-aortic mesenchymal cells. The cellular and molecular components of AGM niche have mainly been investigated using animal models, showing the presumed involvement of BMP and SCF signaling, both with a polarized distribution predominantly to the ventral part of sub-aortic mesenchyme^18, 19^. Interestingly, previous study has described a successful reprogramming from adult mouse ECs to HSCs through transient expression of four TFs combined with an endothelial feeder-dependent induction, emphasizing the role of vascular-niche-derived angiocrine factors in the generation of HSCs^20^. Deciphering the putative interactions that potentially support the specification of HSC-primed hemogenic ECs, the initial step for hematopoietic fate choice, in human embryos is of great importance but has not been achieved.

Single-cell transcriptional profiling has been widely used to decipher the developmental events^21, 22^. It shows unique advantage in the capture of the transient and rare cell populations, such as the emerging HSCs and HSC-primed hemogenic ECs, from invaluable human embryo samples in the present study. Here, we performed both well-based single-cell RNA-sequencing (scRNA-seq) of 528 cells and droplet-based scRNA-seq of 11,440 cells to firstly construct a molecular landscape of HSC generation in human embryos (**Figure S1A**). Precisely understanding the cellular and molecular programs and interactions that underlie the generation of the first HSCs from hemogenic ECs, together with distinguishing HSC-primed hemogenic ECs from the ones related to transient hematopoiesis, will unambitiously shed light on the strategies for the generation of clinically useful HSCs from PSCs.

## Results

### Transcriptional capture of hemogenic ECs in human embryonic dorsal aorta around HSC emergence

We proposed that HSC-primed hemogenic ECs exist at stages around HSC emergence at CS 14^1^, thus should be transcriptionally captured within the aortic structure at CS 13. Therefore, we firstly performed droplet-based scRNA-seq (10X Chromium) with 7-AAD^−^CD235a^−^ cells from the anatomically dissected dorsal aorta in human AGM region at CS 13 (30 dpc). Eight cell clusters featured by the expression of known marker genes were readily recognized and annotated, including EC, arterial EC (AEC), hematopoietic-related cell (Hem), Epithelial cell (Epi) and four distinct mesenchymal cell clusters (**Figures 1A-1C, S1B, S1C and S1E**). As compared to AEC that showed a typical arterial feature such as expression of *GJA5*, *GJA4*, *HEY2*, and *CXCR4*^23–27^, EC cluster exhibited a relatively venous feature such as over-representation of *APLNR*, *NRP2* and *NT5E*^28, 29^ (**Figures 1C and S1B**). The Hem cluster was characterized by the expression of *SPI1* and *SPINK2*^30, 31^ (**Figures 1B and 1C**). Taken together, we successfully captured endothelial and hematopoietic cell populations in the aortic structure, constituting about half of the total cells (48%), together with putative AGM niche cells, mainly involving different types of mesenchymal cells (46%) (**Figure 1D**).

**Figure 1.**
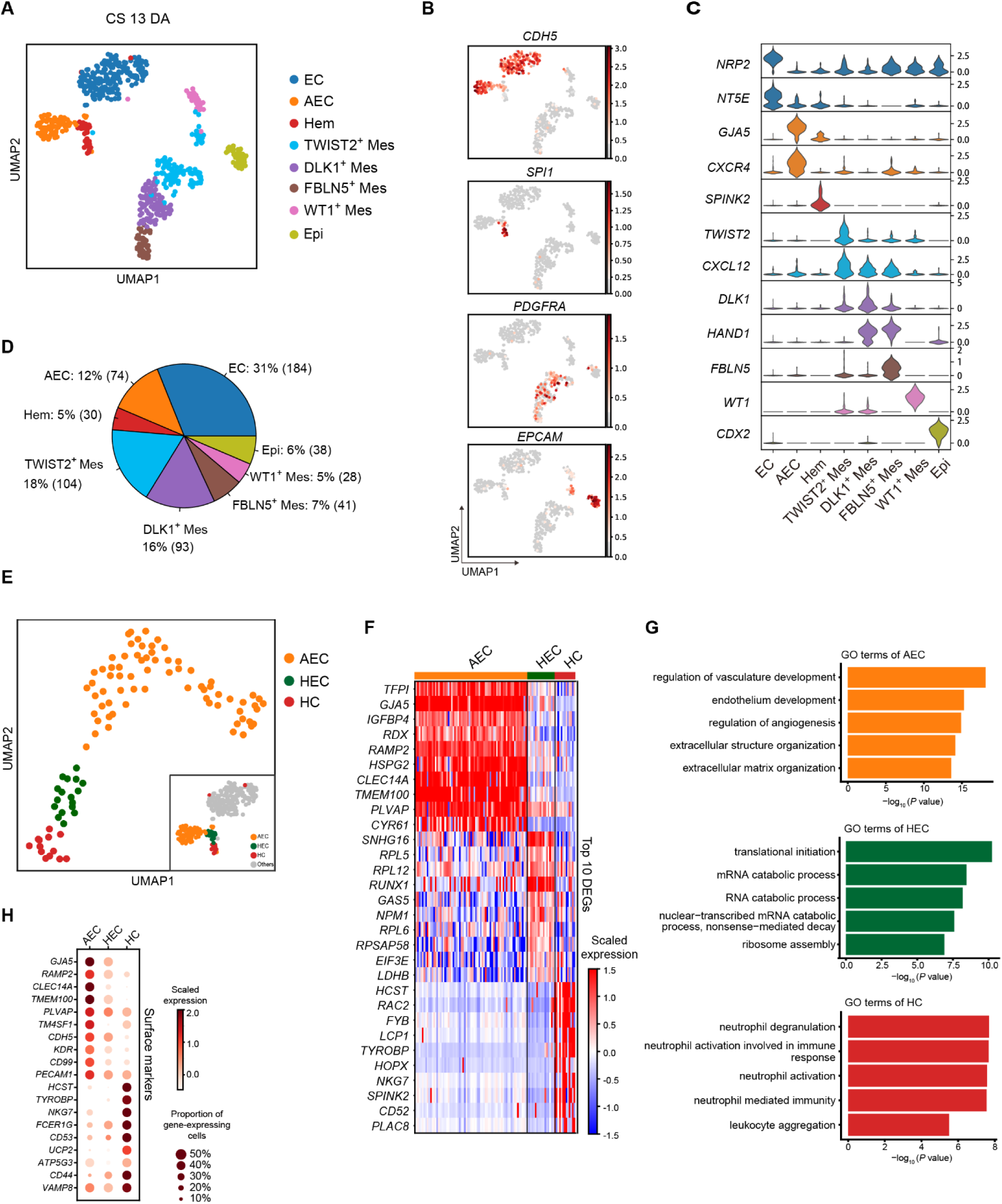
Transcriptomic diversity of cells in human CS 13 dorsal aorta and the capture of hemogenic ECs. **A**. Identification of cell populations in CS 13 DA visualized by UMAP. Each dot represents one cell and colors represent cell clusters as indicated. **B**. UMAP visualization of the expression of curated feature genes for the identification of cell clusters (*CDH5*, *SPI1*, *PDGFRA* and *EPCAM*). **C**. Violin plots showing the expression of feature genes in each cell cluster. Colors represent cell clusters indicated in **Figure 1A**. **D**. Pie charts showing the percentages and absolute numbers of each cell cluster involved in CS 13 DA. **E**. UMAP visualization of AEC, HEC and HC clusters, resulted from sub-dividing cells in AEC and Hem clusters described in **Figure 1A** as indicated in the lower right thumbnail. **F**. Heatmap showing the distinct expression patterns of top 10 differentially expressed genes (DEGs) in AEC, HEC and HC clusters. **G**. The major Gene Ontology biological process (GO:BP) terms in which DEGs are enriched for each cluster. **H**. Dot plots showing the scaled expression level of top 10 significantly differentially expressed surface marker genes in AEC, HEC and HC. Colors represent the scaled expression and size encodes the proportion of gene-expressing cells.

Pearson correlation analysis revealed that AEC showed the highest similarity with Hem cluster, whereas mesenchymal and Epi clusters correlated closely (**Figure S1D**). This finding suggested that the hemogenic activity presumably located within AEC and Hem clusters. To precisely capture the putative hemogenic ECs, AEC and Hem clusters were taken out separately and further divided into three sub-clusters (**Figure 1E**). The largest one was basically corresponding to the previous AEC cluster, thus was still annotated as AEC. Of note, one of the other two sub-clusters showed high expression of *RUNX1* together with endothelial feature, thus was annotated as hemogenic EC (HEC) (**Figures 1E, 1F and S1F**). The other one was named as hematopoietic cell (HC) given the expression of hematopoietic genes *SPN* and *PTPRC* but the lack of endothelial property (**Figures 1E and S1F**). Compared among these three sub-clusters, the major biological processes enriched in AEC were related to extracellular matrix organization and vasculature/endothelium development, in accord with that the dorsal aorta at this stage is undergoing a maturation process^32^ (**Figures 1G and S1C**). In addition to *RUNX1,* a novel long non-coding RNA (lncRNA) - *SNHG16* was found as the most significant differentially expressed genes (DEGs) in HEC (**Figure 1F**). RNA catabolic process was enriched in HEC sub-cluster, also evidenced by the relatively high expression of *RPL5*, *RPL12* and *RPL6*^33^ (**Figures 1F and 1G**).

Next, we computationally screened for the surface markers that might help to enrich and prospectively isolate the hemogenic EC population. Among the three sub-clusters, no surface markers specific for the HEC population were screened out, which was conceivable as the feature of HEC was similar with both AEC and HC clusters. Among the top 10 significantly differentially expressed surface markers between AEC and HC, *CD44* showed relatively more abundant expression in HEC than in AEC, serving as a potential candidate for the enrichment of HEC population (**Figure 1H**).

### Hemogenic ECs in human AGM region exhibited unambiguous arterial feature and were efficiently enriched in phenotypic CD44^+^ ECs

Due to the limited resolution of droplet-based scRNA-seq strategy including 10X Chromium, we subsequently performed well-based scRNA-seq (modified STRT-seq) to more precisely decode the hemogenic ECs in human AGM region at stages shortly before or upon the generation of HSCs (**Figure S1A**). Immunophenotypic CD235a^−^CD45^−^CD34^+^CD44^−^ cells (CD44^−^ ECs) and CD235a^−^CD45^−^CD34^+^CD44^+^ cells (CD44^+^ ECs) were simultaneously sampled with similar cell numbers, although the ratio the latter population took in ECs was at least 10-fold less than the former (**Figure 2A**). Cells were collected from CS 12 (27 dpc) caudal half (CH), CS 13 (29 dpc) and CS 14 (32 dpc) AGM regions of human embryos (**Figure S1A**). An average of 6011 genes were detected in each individual cell and the transcriptional expression of sorting markers basically matched the immunophenotypes (**Figures S2A and S2B**). By unsupervised clustering, the ECs were mainly divided into two populations, largely in line with the immunophenotypes regarding the expression of CD44 (**Figures 2B and S2D**). The cluster composed mainly of CD44^+^ ECs was of arterial feature, with ubiquitous expression of *GJA5*, *GJA4* and *DLL4*, and thus was defined as arterial EC (aEC) (**Figures 2C, 2D and S2E**). The other cluster showed higher expression of venous genes such as *NR2F2* and *NRP2*, and was consequently annotated as venous EC (vEC) (**Figures 2C and 2D**). Of note, hematopoietic gene *RUNX1* was also exhibited in the top 10 over-represented TF genes of aEC population (**Figure 2D**).

**Figure 2.**
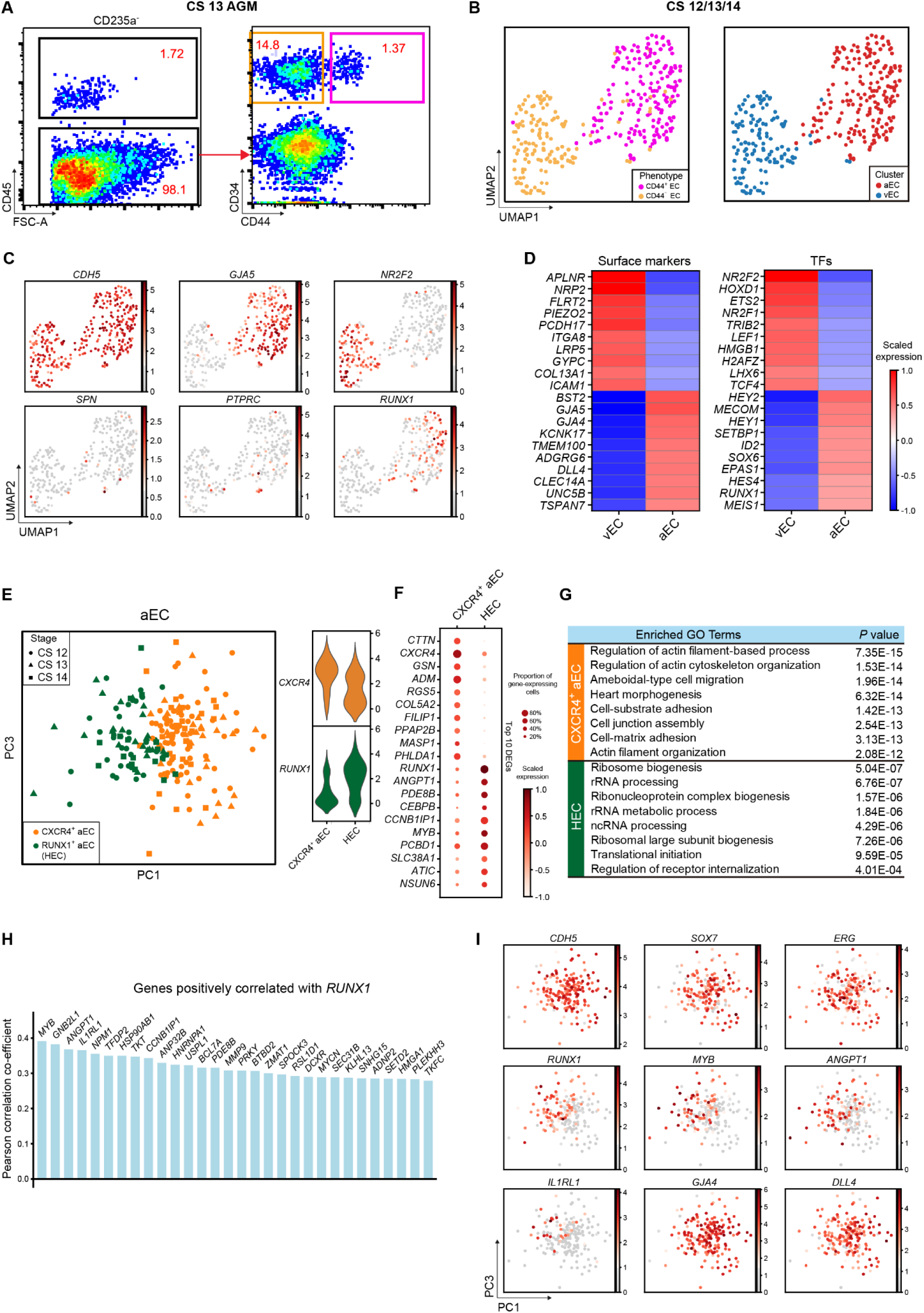
Capture and further analysis of hemogenic ECs in CD44^+^ ECs from CS 12/13/14 embryos. **A**. Sorting strategy of CD44^+^ EC and CD44^−^ EC. **B.** UMAP showing two phenotypically different populations (left panel) can be clustered into arterial EC (aEC) and venous EC (vEC) at the transcriptome level (right panel). Proportion of two clusters is showed in **Figure S2D**. **C**. UMAP plots displaying the expression of hematopoietic and endothelial genes in aEC and vEC clusters. **D**. Heatmaps showing top 10 differentially expressed surface markers and TFs between aEC and vEC clusters. **E**. The aEC cluster is divided into two sub-clusters. The expression of their feature genes is shown on the violin plots on the right panel. **F**. Dot plots showing top 10 DEGs in two sub-clusters. **G**. Enriched GO terms in CXCR4^+^ aEC and HEC, respectively. **H**. Bar plots displaying top genes positively correlated with *RUNX1* in aEC cluster. Genes related to ribosome biogenesis were removed from listed genes. **I**. PCA plot showing expression of endothelial, arterial genes and representative genes from **Figure 2H** in aEC cluster. HEC shares endothelial and arterial feature with CXCR4^+^ aEC. Hematopoietic genes correlated with *RUNX1* enriched in HEC.

Since *RUNX1* was exclusively expressed in a small part of cells in aEC cluster (**Figure 2C**), aEC was further sub-divided in an unsupervised way into two subsets, featured by the expression of *CXCR4* (CXCR4^+^ aEC) and *RUNX1* (HEC), respectively (**Figures 2E, 2F and S2F**). The cellular contributions to each subset were similar among three stages (**Figures S2C and S2F**). Enrichment of pathways including ribosome and translation initiation within HEC was in accord with the role of *RUNX1* in regulating ribosome biogenesis^34^ (**Figures 2G and S2G**). Myb was expressed by HSCs and required for definitive hematopoiesis in mice^35, 36^. Angpt1 was highly expressed by HSCs and might be involved in regulating the regeneration of their niche in murine bone marrow^37^. The respective homologs of these two genes, *MYB* and *ANGPT1*^37, 38^, were included in genes whose expression patterns were positively correlated with that of *RUNX1* (**Figure 2H, Table S1**), and were also enriched in HEC (**Figure 2I**). The expression of *IL1RL1* (the gene encoding receptor for IL33), which was reported co-expressed with *RUNX1* in mouse and human leukemia cells^39^, was also positively correlated with that of *RUNX1* (**Figures 2H and 2I**). Taken together, the HEC cluster, exhibiting a feature of expressing *RUNX1*, *MYB*, *ANGPT1* as well as endothelial genes *CDH5, SOX7* and *ERG* (**Figure 2I**), without apparent expression of hematopoietic surface markers *SPN* and *PTPRC,* was transcriptionally identified as hemogenic EC. These hemogenic ECs were characterized with clear arterial feature represented by the expression of *GJA5*, *GJA4* and *DLL4*^27, 40, 41^ (**Figures 2C, 2D and 2I**). More importantly, considering the physiological cellular constitution, the scarce hemogenic ECs were efficiently enriched in the phenotypic CD44^+^ EC population, by at least 10-fold, in the human AGM region.

### Developmental path from arterial ECs via HSC-primed hemogenic ECs to HSPCs in human AGM region

Functionally, the most efficient marker combination reported to enrich the emerging HSCs in human AGM region is CD34^+^CD45^+3^. Although other markers have been described, the enrichment is not further improved^3^. In order to cover the rare HSCs, here we performed well-based scRNA-seq with sorted CD235a^−^CD45^+^CD34^+^ cells, which were proven to contain most, if not all, functional human AGM HSCs by xenotransplantations^3^, from the dorsal aorta in human AGM region at CS 15 (**Figure 3A**). By unsupervised clustering, the immunophenotypic CD235a^−^CD45^+^CD34^+^ population was divided into five clusters (**Figure 3B**). Among them, two clusters exhibited feature of hematopoietic differentiation, with one towards myeloid lineage represented by the obvious expression of *GATA1* and *ITGA2B* and the other having the sign of lymphoid lineage potential evidenced by the expression of *IL7R and LEF1*^42, 43^ (**Figures 3C and S3A**). The remaining three clusters with stemness signatures (*FGD5* and *HLF*) were thus recognized as HSPCs, constituting about 75.4% of the immunophenotypic CD235a^−^CD45^+^CD34^+^ population^44, 45^ (**Figures 3B and 3C**). Notably, as a marker previously recognized for identifying functional long-term HSCs in mouse bone marrow, *FGD5* was among the top 10 DEGs highly expressed in HSPCs when compared to the two differentiated populations^44^ (**Figures 3C and 3D**). *HLF*, as reported to be expressed in HSCs in multiple mouse embryonic hematopoietic sites and might maintain their quiescence^45, 46^, was included in top 10 TFs enriched in HSPC clusters (**Figures 3C and S3A**). The results further validated our identification of HSPCs in dorsal aorta.

**Figure 3.**
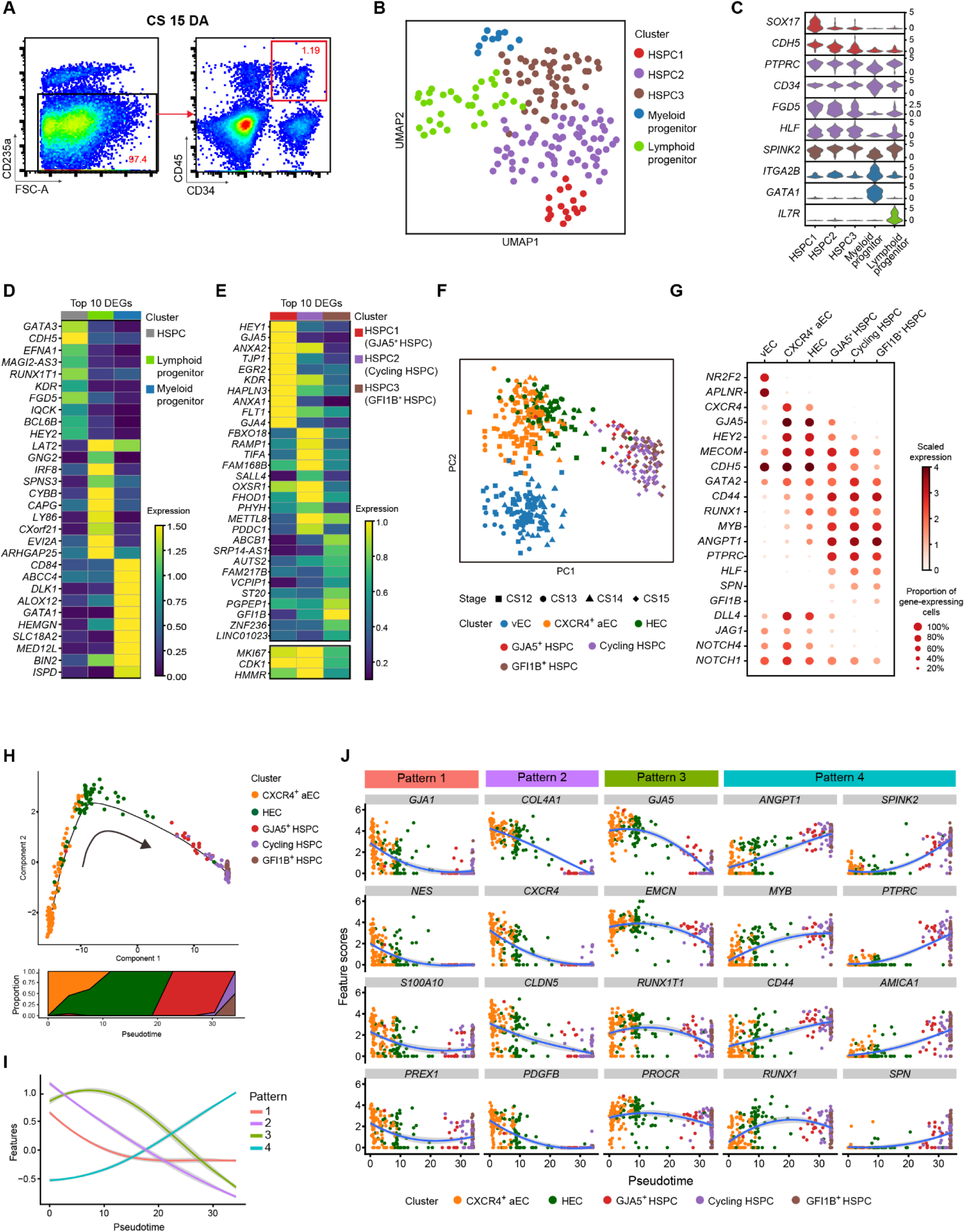
The developmental path from arterial ECs via HSC-primed hemogenic ECs to HSPCs in AGM region. **A**. Sorting strategy of CD235a^−^CD45^+^CD34^+^ hematopoietic progenitors in CS 15 dorsal aorta. **B**. Identities of five cell populations in CS 15 dorsal aorta visualized by UMAP. **C**. Violin plots showing the expression of feature genes in each cell cluster. **D**. Heatmap showing the top 10 DEGs expressed in HSPC (HSPC1/2/3), Myeloid progenitor and Lymphoid progenitor clusters. **E**. Heatmap showing the distinct expression patterns of top 10 DEGs in HSPC1 (GJA5^+^ HSPC), HSPC2 (Cycling HSPC) and HSPC3 (GFI1B^+^ HSPC) clusters. **F**. PCA plot of vEC, CXCR4^+^ aEC, HEC, GJA5^+^ HSPC, Cycling HSPC and GFI1B^+^ HSPC. **G**. Dot plots showing the scaled expression level of feature genes in six cell clusters mentioned in **Figure 3F**. Expression of Notch signaling pathway genes is shown at the bottom. **H**. Pseudotime analysis of two aEC sub-clusters (CXCR4^+^ aEC and HEC) from CS 12/13/14 and three HSPC clusters (GJA5^+^ HSPC, Cycling HSPC and GFI1B^+^ HSPC) from CS 15 indicates the developmental trajectory by Monocle 2. Dynamic changes of proportion of clusters are shown on the bottom panel. **I**. Four distinct gene expression patterns along the pseudotime axis. **J**. Expression of representative genes of each pattern along endothelial-to-hematopoietic transition (EHT) process.

Three HSPC clusters showed distinct features. The one farthest from the lineage-primed clusters, which should be developmentally the most upstream population within CD45^+^CD34^+^ cells (**Figure 3B**), manifested higher endothelial (*KDR* and *FLT1*) and arterial (*HEY1*, *GJA5* and *GJA4*) features as compared to the other two HSPC populations, thus was annotated as GJA5^+^ HSPC (**Figure 3E**). The finding suggested arterial-featured hemogenic ECs as the source of the first HSCs in human AGM region. The most significant difference between the other two HSPC clusters was the cell cycle status, and the one with highly proliferative property was consequently annotated as Cycling HSPC (**Figure S3C**). The other one was annotated as GFI1B^+^ HSPC given its highest expression of *GFI1B* among the three HSPC clusters (**Figure 3E**). As a direct target of *Runx1*, *Gfi1b* is a marker whose expression can be used in enriching mouse AGM HSCs with long-term repopulating capacity^47^.

To investigate the molecular events during the emergence of HSCs, we pooled together the datasets of relevant cell populations from modified STRT-seq data, including vEC, CXCR4^+^ aEC and HEC in CS 12 (CH), CS 13 and CS 14 of AGM region (**Figures 2B and 2E**), and the three HSPC populations in CS 15 for further analysis. Dimension reduction was performed by principal component analysis (PCA) (**Figure 3F**). Of note, principal component (PC) 1 captured the differences between endothelial and hematopoietic properties, showing HEC and GJA5^+^ HSPC located adjacent to the middle part of the axis (**Figures 3F and S3B**). PC2 positive direction represented the arterial feature, exhibiting the gradual down-regulation along with hematopoietic specification of HSPCs from HEC (**Figures 3F and S3B**). The result was in line with previous reports in mouse embryos showing reciprocal relationship of arterial genes expression and hematopoietic fate acquisition^48, 49^. In contrast to the lack of the expression of venous feature genes, including *NR2F2* and *APLNR*, HSPCs expressed different levels of canonical arterial genes, such as *GJA5*, *HEY2,* and *MECOM*^50^, indicative of the arterial EC origin of HSCs (**Figure 3G**). The expression patterns of *CD44*, *RUNX1*, *MYB* and *ANGPT1* were similar, being seldom expressed in vEC, gradually increased from CXCR4^+^ aEC via HEC to HSPCs, with obvious expression in all HSPC clusters (**Figure 3G**). Of note, *HLF*, *SPN* and *GFI1B* were exclusively expressed in three HSPC clusters^45^ (**Figure 3G**). In accord with their involvement in arterial specification^48^, several genes of NOTCH signaling pathway, including *DLL4*, *JAG1*, *NOTCH1* and *NOTCH4*^14^, were most highly expressed in CRCR4^+^ aEC, and gradually down-regulated along the specification via HEC towards HSPCs (**Figure 3G**). We also witnessed the difference in cell cycle status among distinct clusters. CXCR4^+^ aEC and HEC showed a relatively quiescent status as compared to vEC, with a dramatically increased proliferation in HSPCs (**Figure S3C**). These findings suggested that once the hematopoietic fate is acquired, HSPCs may rapidly expand *in situ* before differentiation.

Monocle analysis revealed the developmental trajectory of endothelial-to-hematopoietic transition in human AGM region, clearly showing the path from CXCR4^+^ aEC via HEC, GJA5^+^ HSPC, to Cycling and GFI1B^+^ HSPC clusters (**Figures 3H and S3D**). Genes that were significantly differentially expressed along the pseudotime axis were subjected to k-means clustering analysis, resulting in four distinct gene expression patterns (**Figures 3I, 3J, S3E and S3F, Table S2**). Genes grouped in pattern 1 highly expressed only in CXCR4^+^ aEC while rapidly decreased in HEC and HSPCs, represented by *GJA1*, *NES*, *S100A10* and *PREX1*, were related to the regulation of cytoskeleton organization. Genes in both pattern 2 and pattern 3 were related to angiogenesis, EC migration and cell junction organization, with the former simultaneously sharing the biological terms with pattern 1 (**Figure S3F**). Genes in pattern 2 showed gradual decrease in expression accompanied with hematopoietic specification, represented by *COL4A1*, *CXCR4*, *CLDN5* and *PDGFB* (**Figures 3J and S3E**). Genes in pattern 3 maintained or even slightly increased expression level upon the initial step of hemogenic fate choice of arterial ECs, while being down-regulated in HSPCs, represented by *GJA5, EMCN, RUNX1T1* and *PROCR* (**Figures 3J and S3E**). Some of these genes were validly studied in HSCs. In mice, functional HSCs in E11.5 AGM region are exclusively within Emcn^+^ cells based on CD45^+^CD41^+^ phenotype^51^. *PROCR* is reported to efficiently enrich HSCs in human fetal liver^52^ and its homolog has been proven to be a marker for type I and type II pre-HSCs in mice^53^. *RUNX1T1*, whose homolog in mice, with other five TFs, could impart multi-lineage potential to lineage-primed hematopoietic progenitors in reprograming research, implying its function in initiating the molecular program towards multi-potential HSPCs^54^. Genes in pattern 4 were those up-regulated in HSPCs, including the ones already highly expressed in HEC as compared to CXCR4^+^ aEC, such as *ANGPT1*, *MYB*, *CD44* and *RUNX1* (**Figures 3J and S3E**), and also those specifically expressed in HSPCs, such as *SPINK2*, *PTPRC*, *AMICA1 and SPN* (**Figures 3J and S3E**). Of note, *SPINK2* has been reported to be expressed in HSCs in human umbilical cord blood^31^.

### A distinct hemogenic EC population lacking arterial feature existed prior to HSC-primed hemogenic ECs in human embryos

As the HSC-primed hemogenic ECs in human embryos could be detected at CS 12, to explore whether hemogenic ECs existed at stages much earlier, cells from CS 10 body part and CS 11 CH of embryo proper were collected for droplet-based scRNA-seq and analyzed together. The strategy applied to define different populations was the same as that used in analysis for the data from CS 13 DA (**Figures 1A-1C**). For clearer presentation, populations from CS 10/11 were prefixed with “early” while those from CS 13 were prefixed with “late” (**Table S3**). Endothelial population (Endo), featured by *CDH5* and *SOX7* expression, and hematopoietic population with *RUNX1* expression much adjacent to Endo (Hema) were picked out from CS 10/11 dataset and re-analyzed in searching of hemogenic ECs (**Figures S4A and S4B**). Another hematopoietic population showing clear macrophage identity evidenced by *C1QC* and *CD68* expression was not included (**Figures S4A and S4B**).

Three EC populations and two hematopoietic populations were readily recognized by their distinct feature genes. They were vascular ECs with primitive and venous features (early EC), cardiac ECs (early cEC), arterial ECs (early AEC), megakaryocytic progenitors (early Mega) and erythroid progenitors (early Ery), respectively (**Figures 4A-4C and S4C**). A putative hemogenic EC population, a part of which met the strict definition of hemogenic EC by expressing endothelial genes (*CDH5* and *ERG*) and *RUNX1* but lacking expression of hematopoietic marker genes (*ITGA2B, SPN* and *PTPRC*), was identified and annotated as early HEC (**Figures 4A and 4B**). The early HEC located nearer to EC populations than to hematopoietic populations in Uniform Manifold Approximation and Projection (UMAP) (**Figure 4A**), implying their endothelial identity. Specifically, they highly expressed *CD44* and *SPI1* as compared to other endothelial and hematopoietic populations (**Figure 4C**). To strictly comply with the definition of hemogenic EC, a part of cells within early EC cluster expressing specific hematopoietic markers (*ITGA2B, SPN* or *PTPRC*), which located further away from EC populations, were excluded from early HEC cluster and separately named as early HC in the subsequent analysis.

**Figure 4.**
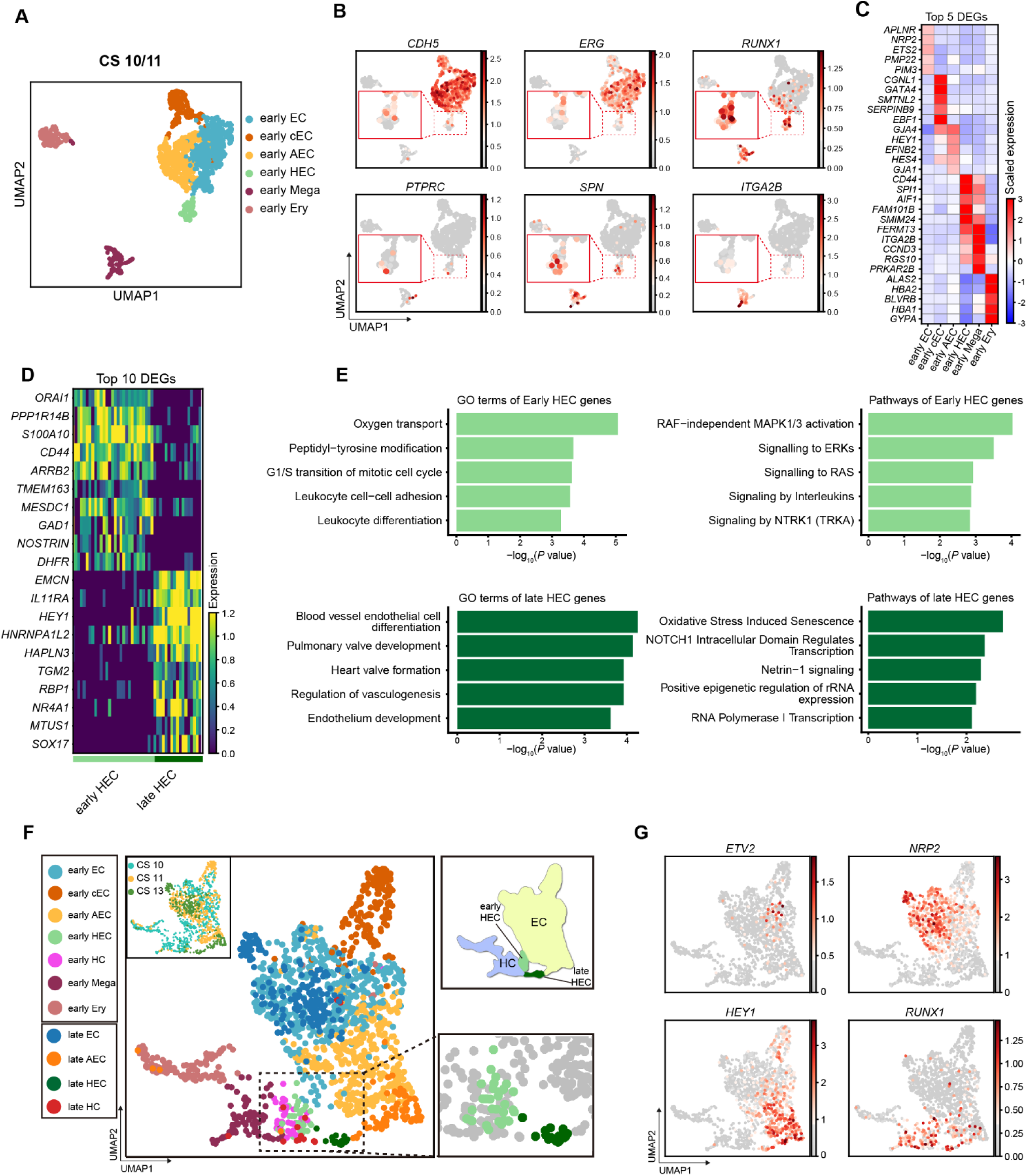
Different feature and origin of the early and late hemogenic ECs. **A**. UMAP showing early endothelial and hematopoietic populations in CS 10 and CS 11 of human embryos. Early HEC lies more closely to endothelial cells. **B**. Expression patterns of typical endothelial and hematopoietic genes in early populations (CS 10 and CS11). **C**. Heatmap showing the expression of top 5 DEGs enriched in distinct populations. **D**. The expression of top 10 DEGs in early and late HEC. **E**. GO:BP terms and pathways enriched in early HEC (upper panel) and late HEC (lower panel). **F**. UMAP of endothelial and hematopoietic cells from early (CS 10 and CS 11) and late (CS 13) stages. **G**. Expression patterns of genes that can depict a developmental pathway in **Figure 4F**.

Different molecular features were found in early and late HEC. *ORAI1*, *PPP1R14B* and *S100A10* were among the top of DEGs expressed in early HEC (**Figure 4D, Table S4**). Expression of genes representing an arterial feature including *HEY1* and *SOX17* was enriched in late HEC (**Figure 4D**). *IL11RA* was also up-regulated in late HEC, accordant with the multiple biological function of its ligand IL11 in lymphohematopoietic cells^55^ (**Figure 4D**). Gene ontology (GO) terms related to oxygen transport, leukocyte cell-cell adhesion and differentiation were enriched in early HEC (**Figure 4E**). In comparison, those related to endothelial cell differentiation, development and vasculogenesis were enriched in late HEC, indicating a more mature endothelial feature of them (**Figure 4E**). Pathways related to NOTCH1 intracellular domain, positive epigenetic regulation of rRNA expression and RNA polymerase I transcription were enriched in late HEC, revealing a Notch-dependent arterial property together with an activated transcriptional and translational feature of late HEC (**Figure 4E**).

Since HSC emergence occurs almost 10 days later than CS 10 (22-23 dpc)^1^, it would be very likely that hemogenic ECs existing at CS 10/11 (early HEC) and CS13 (late HEC) had different derivations and represented endothelial precursors of different hematopoietic populations. Thus, endothelial, hemogenic and hematopoietic populations collected from CS 10/11 (**Figure 4A**) were pooled with their counterparts from CS 13 embryo (**Figure 1A**) for further analysis (**Figure 4F**). Dimension reduction showed that a small population expressing the primitive EC feature gene *ETV2*^56^, which was predominantly sampled from CS 10/11, was localized in the center of EC populations, with one direction towards venous feature and the other towards arterial feature specification (**Figures 4F and 4G**). In contrast to late HEC that was specified from late AEC with the strongest arterial feature, early HEC was segregated from the ECs showing ambiguous arteriovenous feature, indicating their generation did not transit through an arterial stage (**Figures 4F and 4G**). Therefore, two distinct hemogenic EC populations formed two bridges connecting EC populations with hematopoietic populations (**Figure 4F, upper right panel**). Notably, few cells in early HEC were from CS 11 (3 cells) while plenty of them (26 cells) from CS 10, albeit the numbers of endothelial populations sampled from both stages were comparable (**Figures 4A and S4D**). The temporal gap between the detection of two hemogenic EC populations excluded the possibility that late HEC might be developed from early HEC, and more likely, they developed independently and gave rise to different hematopoietic progenies, which were distinguished at least by whether they contain HSCs.

### Computational analysis of the heterologous cellular interactions for the development of HSC-primed hemogenic ECs in human embryos

The ventral wall of dorsal aorta is the site detecting the first HSCs in human embryos^3^. However, the cellular constitution and molecular basis of the dorsal aortic niche remains unknown. Using the scRNA-seq dataset sampled from the anatomically dissected dorsal aorta in AGM region at CS 13, we investigated the previously unknown cellular components and potential cellular interactions acting on the hemogenic ECs, the development of which serves as the initiation of the cell fate specification towards HSCs. In addition to endothelial and hematopoietic populations, five cell populations, including epithelial cell (Epi) and four molecularly different mesenchymal cell populations (Mes), were identified (**Figure 1A**). Combined with feature genes and GO terms, these populations were readily recognized (**Figures 1C, S1B and S1C**). Epithelial cells, featured by *EPCAM* and *CDX2* expression (**Figures 1B and 1C**), were likely the cells from developing gastrointestinal tract adjacent to the ventral wall of dorsal aorta^57, 58^. WT1^+^ Mes was probably from the mesonephros tissues nearby the dorsal aorta^59^. The remaining three mesenchymal populations extensively expressed *PDGFRA* (**Figure 1B**), which marks the mesenchyme around dorsal aorta^60^. Of note, both *Dlk1* and *Fbln5* are expressed in aortic smooth muscle cells in mice^61, 62^. The TWIST2^+^ Mes seemed to have multiple differentiation capacity, and might be the undifferentiated mesenchymal progenitors^63^.

The cellular interactions were represented by the simultaneous expression of heterologous ligand-receptor gene pairs. Regarding mesenchymal and epithelial populations as potential niche components, only ligands expressed by these populations and receptors expressed by HEC were considered by us (**Figure 5A**). Among a total of 128heterologous ligand-receptor pairs detected, cytokine-cytokine receptor interaction, axon guidance, endocytosis, HEDGEHOG signaling pathway, and TGF-beta signaling pathway were the pathways predominantly involved (**Figure 5B**). Those related to NOTCH signaling pathway, focal adhesion, MAPK signaling pathway, dorso-ventral axis formation and MTOR signaling pathway were also recurrent (**Figure 5B**). The ligand-receptor gene pairs exhibited different expression patterns among distinct stromal populations when coupled with HEC (**Figures 5C and S5A**). Among three sub-aortic mesenchymal populations, DLK1^+^ Mes and FBLN5^+^ Mes, paired with HEC, showed relatively similar expression patterns of heterologous ligand-receptor gene pairs, while TWIST2^+^ Mes showed only a few interactions with HEC (**Figure 5C**). DLK1-NOTCH1 were presented in DLK1^+^ Mes and FBLN5^+^ Mes, when coupled with HEC (**Figures 5D and S5A**). More specifically, SPP1-CD44, WNT2B-FZD4 and IL11-IL11RA pairs were presented in TWIST2^+^ Mes, DLK1^+^ Mes and FBLN5^+^ Mes, when coupled with HEC, respectively (**Figure 5D**). BMP4 involved pairs were shared by WT1^+^ Mes, DLK1^+^ Mes and FBLN5^+^ Mes, including BMP4-BMPR2-BMPR1A, BMP4-AVR2A-BMPR1A and BMP4-AVR2B-BMPR1A (**Figures 5D and S5A**), accordant with previous report that BMP4 is expressed in the ventral sub-aortic mesenchyme of human embryos^64^ and the suggested role of Bmp4 in HSC generation in mice^65, 66^.

**Figure 5.**
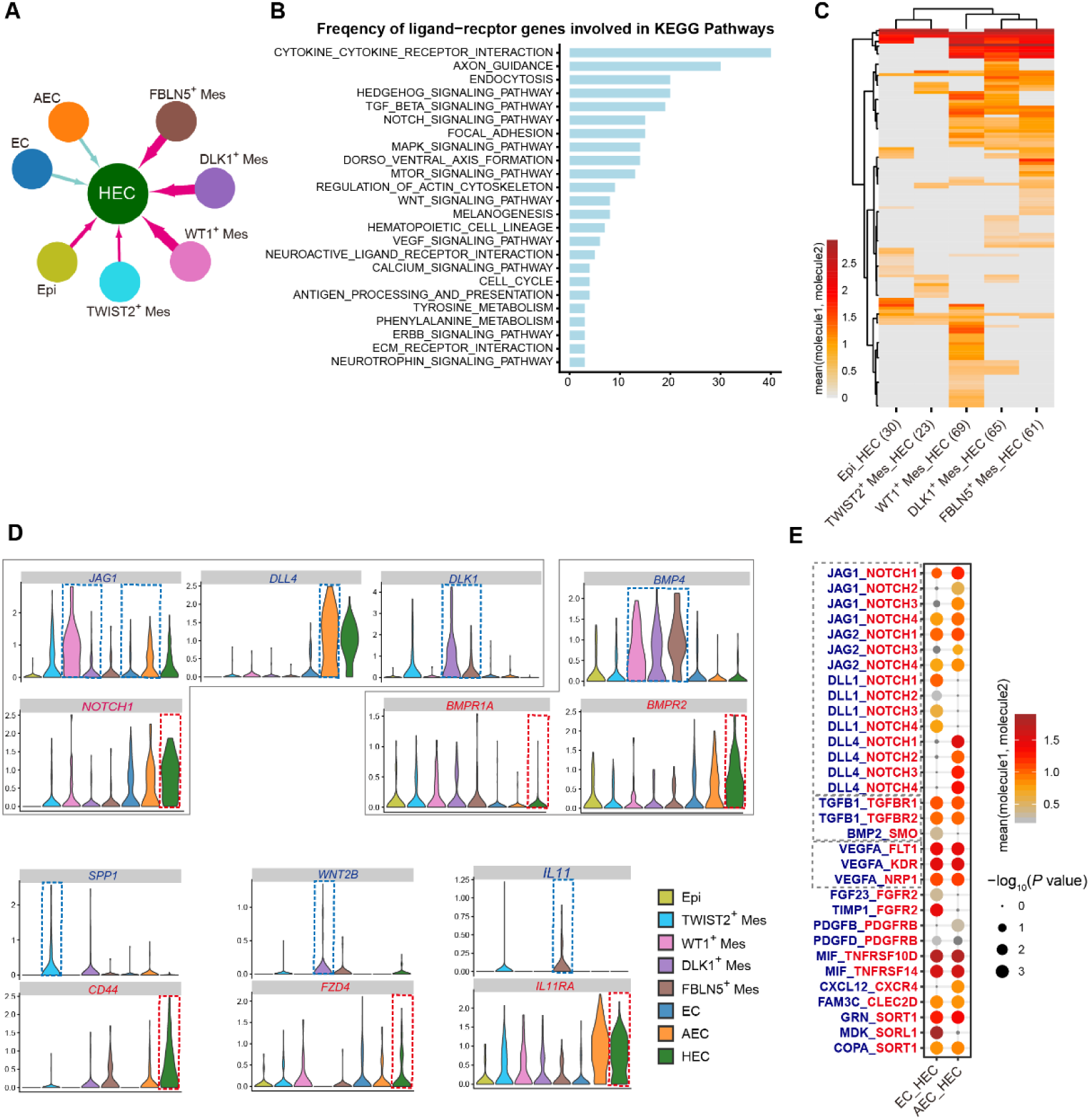
Computational analysis of the heterologous cellular interactions for the development of HSC-primed hemogenic ECs. **A**. The schematic diagram indicates the ligand-receptor interactions between HEC and other clusters including EC, AEC, epithelial and four mesenchymal cell clusters. **B**. Frequency of ligand-receptors involved in KEGG pathways. **C**. Heatmap showing the ligand-receptor pairs that exhibit different expression patterns among distinct stromal clusters when coupled with HEC. **D**. The expression of ligand and its receptor genes of indicated pairs, including DLK1/JAG1/JAG2/DLL4-NOTCH1, SPP1-CD44, WNT2B-FZD4 and IL11-IL11RA. **E**. Dot plots showing the ligand-receptor pairs between either EC or AEC and HEC.

Considering the special architecture of endothelial layer in dorsal aorta, interactions among endothelial populations should most likely depend on molecules that can act remotely. Thus, cytokines and chemokines were taken with Notch signaling into consideration in cell-cell interaction analysis. The interactions between either EC or AEC and HEC included NOTCH-related (JAG-NOTCH and DLL-NOTCH), TGFB-related (TGFB1-TGFBR1, TGFB1-TGFBR2 and BMP2-SMO) and VEGF-related (VEGFA-FLT1, VEGFA-KDR and VEGFA-NRP1) (**Figure 5E**).These interactions have been widely studied in vertebrate embryonic hematopoiesis and for *in vitro* generation of HSCs from PSC^67, 68^. However, their functional involvement in human HSC generation needs further investigations.

## Discussion

In the present study with the precious human embryonic tissues, we constructed for the first time a genome-scale gene expression landscape for HSC generation in the AGM region of human embryos, in which we focused specifically on the cell populations and molecular events involved in the endothelial-to-hematopoietic transition. We transcriptomically identified the HSC-primed hemogenic ECs, meeting the widely acknowledged criteria for hemogenic ECs that simultaneously express core feature genes of ECs (such as *CDH5* and *SOX7*) and critical hematopoietic TFs (such as *RUNX1* and *MYB*) but yet not specific surface markers of hematopoietic cells (including *PTPRC* and *SPN*)^9, 69^ (**Figures 2C and 2I**). The hemogenic ECs exhibited a much more intimate relationship with vascular ECs than with the functionally validated HSPCs showing an immunophenotype of CD45^+^CD34^+^ (**Figure 3F**), confirming their endothelial identity and origin. The finding is in line with the notion in the mouse embryos that non-hemogenic ECs and hemogenic ECs in AGM region show great similarity in transcriptome^70^ (**Figures 3F and 3G**). Of note, *RUNX1* was ranked as the top one DEG when the EC population with an arterial feature in AGM region was further sub-divided in an unsupervised way (**Figure 2F**). Together with the specific GO terms and enriched pathways (**Figures 2G and S2G**), the transcriptomically identified hemogenic ECs here should be recognized as the earliest cell population specified to choose a hematopoietic fate, which has not been uncovered in human embryos.

We revealed that the HSPCs in CS 15 AGM region showed certain levels of arterial feature but nearly absent of venous feature (**Figure 3G**), in line with the reports in the mouse embryos that functional type I and type II pre-HSCs in the AGM region express evident arterial markers but much lower levels of venous markers^53^. The data suggested an arterial EC origin of emerging HSCs in human, in accord with the apparent arterial feature of the hemogenic ECs we transcriptomically identified in the CS 12-14 AGM region (**Figure 3G**), which is previously unknown in human embryos. Concerning the *in vitro* differentiation system of human PSCs, the arterial-specific markers DLL4 and CXCR4 can be used to identify hemogenic ECs with lympho-myeloid potential^14^. Interestingly, we found that the hemogenic EC sub-population showed much lower *CXCR4* expression than the other one with comparable arterial feature in the AGM (**Figure 2F**). Thus, albeit *CXCR4* is regarded as an arterial gene^17, 71^, CXCR4^+^ ECs represented a proportion but not all of arterial ECs, showing much less hemogenic characteristics. Therefore, it can be explained that CXCR4^+^ cells generated from human PSCs *in vitro* lack direct hematopoietic potential^71^. Our finding emphasizes the importance of discriminating cell identity transcriptomically at single-cell resolution and further suggests that *CXCR4* expression might be rapidly down-regulated upon the hemogenic specification in the arterial ECs.

Transcriptionally, the homing receptor *CD44* was expressed in a subset of ECs with arterial feature but seldom in venous ECs of AGM region in human embryos (**Figures 2B and 2C**). Importantly, almost all the cells with hemogenic characteristics were enriched in the immunophenotypic CD44^+^ EC population, and the expression of *CD44* was gradually increased along with hematopoietic specification (**Figure 3G**). The results are in line with the histological finding that CD44 is expressed in the IAHC cells, as well as the nearby ECs lining the inner layer of the dorsal aorta of human embryos at the stages similar to ours^72^. Therefore, together with CD44 labeling, which constituted an average 7.4% of CD34^+^CD45^−^ ECs in AGM region, the hemogenic ECs in human embryos are enriched, for the first time, more than 10-fold as compared to that by pan-EC markers. Combining flow cytometry with computational identification, the transcriptomically defined hemogenic ECs constituted about 1/30-40 of the ECs in AGM region. Previous study has reported the frequency of blood-forming ECs in human AGM at about 1/150 of CD34^+^CD45^−^ cells at 30 dpc (CS 13) evaluated by an *in vitro* co-culture system^73^. It remains to be determined that if all the transcriptomically defined hemogenic ECs in the present study are functional, especially with the improvement of the culture system and even *ex vivo* functional evaluation.

In an effort to determine the earliest time-point detecting the intra-embryonic hemogenic ECs in human embryos, we unexpectedly revealed two temporally and molecularly distinct hemogenic EC populations, with the earlier one lacking obvious arterial feature (**Figure 4C**). It has been proven that distinct waves of hematopoiesis occur sequentially in model organisms including mouse and zebrafish embryos, with both transient definitive non-HSC hematopoiesis and definitive HSCs generated from hemogenic ECs^8, 73^. We proposed that the previously undefined two distinct intra-embryonic hemogenic EC populations might correspond to the putative two definitive waves of hematopoiesis in human embryos (**Figure 4F**). The functional difference between these two transcriptomically identified intra-embryonic hemogenic EC populations in human embryos needs further investigations. It is to be determined that whether two hemogenic EC populations we transcriptomically identified have the common ancestor, such as the immature hemogenic ECs proposed in the PSC differentiation system^14, 17^. Much likely, along the arterial specification path from primordial ECs, two cohorts of the intra-embryonic hemogenic ECs are sequentially segregated out, with one before and the other after arterial fate settling. It is extremely pivotal to discriminate the initial steps for different types of hematopoiesis during human embryogenesis, which are presumably related to quite distinct self-renewal and differentiation capacities and physiological functions. More importantly, this will provide crucial clues for the *in vitro* blood regeneration studies.

## Materials and Methods

### Ethics statement and sample collection

Healthy human embryonic samples were obtained with elective medical termination of pregnancy in Affiliated Hospital of Academy of Military Medical Sciences (the Fifth Medical Center of the PLA General Hospital). All experiments were performed in accordance with protocols approved by the Ethics Committee of the Affiliated Hospital of Academy of Military Medical Sciences (ky-2017-3-5), and local and state ethical guidelines and principles. The written informed consent was obtained before sample collection. Carnegie stages (CS) were used ^74, 75^ to determine the stages of embryos according to crown-rump length (CRL) measurement and number of somite pairs. Samples used in this study were from CS 10 (23 dpc), CS 11 (24 dpc), CS 12 (27 dpc), CS 13 (29 and 30 dpc), CS 14 (32 dpc) and CS 15 (36 dpc) embryos (**Figure S1A**).

### Preparation of single-cell suspensions

Human embryonic body, caudal half and AGM region were isolated and transferred to IMDM medium (Gibco, 12440053) containing 10% Fetal Bovine Serum (HyClone, SH30070.03) on ice. AGM region was washed by PBS and transferred to pre-warmed digestion medium containing 0.1 g/mL Collagenase I (Sigma, C2674), which was shaken vigorously for 30 seconds and further incubated at 37 °C for about 30 minutes in incubator with gentle shaking every 5 minutes to release cells. IMDM medium containing 10% Fetal Bovine Serum was added to terminate digestion. Cells were then collected by centrifuging at 350 g for 6 minutes, and resuspended in FACS sorting buffer (1× PBS with 1% BSA) for subsequent staining.

### Fluorescence activated cell sorting (FACS) and single-cell RNA-sequencing

Cells were stained in FACS sorting buffer with specific antibodies for 30 min at 4 °C. The following antibodies were used: BV421-conjugated anti-human CD45 (BD Biosciences, 563879), FITC-conjugated anti-human CD34 (BD Biosciences, 555821), APC-Cy7-conjugated anti-human CD235a (BioLegend, 349116), BV605-conjugated anti-human CD44 (BD Biosciences, 562991) and PerCP-Cy5.5-conjugated 7-AAD (eBioscience, 00-6993-50). After staining, cells were washed once and resuspended in FACS sorting buffer. Cells were sorted on BD FACS Aria II. Pre-gating was first done for live cells based on a 7-AAD staining. Data analysis was performed using FlowJo V10 software (https://www.flowjo.com).

### Single cell library construction

Droplet-based scRNA-seq datasets were produced with a Chromium system (10X Genomics, PN120263) following manufacture’s instruction. For well-based scRNA-seq, sorted single cells in good condition were picked into lysis by mouth pipetting, and the scRNA-seq libraries were constructed based on STRT-seq with some modifications ^76–78^. cDNAs were synthesized using sample-specific 25-nt oligo-dT primer containing 8-nt barcode (TCAGACGTGTGCTCTTCCGATCT-XXXXXXXX-DDDDDDDD-T25, X representing sample-specific barcode while D standing for unique molecular identifiers (UMI), shown in **Table S5**) and template switch oligo (TSO) primer for template switching^79–81^. After reverse transcription and second-strand cDNA synthesis, the cDNAs were amplified by 16 cycles of PCR using ISPCR primer and 3’ Anchor primer (see **Table S5**). Samples were pooled and purified using Agencourt AMPure XP beads (Beckman, A63882). 4 cycles of PCR were performed to introduce index sequence (shown in **Table S5**) and subsequently, 400 ng cDNAs were fragmented to around 300 bp by Covaris S2. After being incubated with Dynabeads MyOneTM Streptavidin C1 beads (Thermo Fisher, 65002) for 1 hour at room temperature, cDNA Libraries were generated using KAPA Hyper Prep Kit (Kapa Biosystems, kk8505). After adaptor ligation, the libraries were amplified by 8 cycles of PCR using QP2 primer and short universal primer (shown in **Table S5**). The libraries were sequenced on Illumina Hiseq X Ten platform in 150 bp pair-ended manner (sequenced by Novogene).

### Processing single-cell RNA-seq data

Sequencing data from 10X genomics was processed with CellRanger software package (version 2.1.0) with default mapping arguments. For modified STRT-seq data, raw reads were first split for each cell by specific barcode sequence attached in Read 2. The TSO sequence and polyA tail sequence were trimmed for the corresponding Read 1 after UMI information was aligned to it. Reads with adapter contaminants or low-quality bases (N > 10%) were discarded. Subsequently, we aligned the stripped Read 1 sequences to hg19 human transcriptome (UCSC) using Hisat2 (version 2.10)^82^. Uniquely mapped reads were counted by HTSeq package^83^ and grouped by the cell-specific barcodes. Duplicated transcripts were removed based on the UMI information for each gene. Finally, for each individual cell, the copy number of transcripts of a given gene was the number of the distinct UMIs of that gene.

### Quality control

Then quality control was performed to filter low-quality cells. For 10X-derived datasets, only cells with more than 1,000 genes less than 1,000,000 UMIs were retained for downstream analysis. For modified STRT-seq datasets, UMI values for each cell were grouped in an expression matrix, cells were retained when more than 2,000 genes and 10,000 UMIs were detected.

### Dimension reduction and clustering

The Seurat (version 2.3.4) ^84^ implemented in R (version 3.4) was applied to reduce the dimension of CS 13 10X-derived datasets. UMI count matrix was applied a logarithmic transformation with the scale factor 10,000. High variable genes (HVGs) were calculated using FindVariableGenes function with default parameters except for “x.low.cutoff” = 0.0125. PCA was performed using HVGs, and significant PCs were selected to perform dimension reduction and clustering. Cells were projected in 2D space using UMAP with default parameters. Using graph-based clustering function Findcluster with default parameters except for “resolution” = 0.6, we divided the 592 cells into 8 transcriptionally similar clusters (**Figure 1A**). For CS 12/13/14 modified STRT-seq data, the Normalization scale factor was set to 100,000. The PCA was performed using HVGs with parameter “x.low.cutoff” = 0.2, and then UMAP analysis with parameters “n_neighbors” = 10, “min_dist” = 0.3 and clustering with parameter “resolution” =0.8, resulting in 2 endothelial clusters (**Figure 2B**). For Hemogenic EC analysis, we recalculated HVGs and performed PCA on the aEC subset, and PC1 and PC3 captured most of the variation between populations. CS 15 modified STRT-seq data was subjected to Scanpy (version 1.4.2) ^85^, 5 clusters were identified using Louvain clustering with 5 nearest neighbors and first 20 PCs, and UMAP was used for dimension reduction and visualization of gene expression.

### Integrated analysis of CS 10/11/13 10X-derived data

To account for batch differences among CS 10, CS 11 and CS 13 10X-derived datasets, we used the Batch balanced KNN (BBKNN), a batch effect removal tool in Scanpy package. BBKNN actively combats technical artifacts by splitting data into batches and finding a smaller number of neighbors for each cell within each of the groups, which helps create connections between analogous cells in different batches without altering the counts or PCA space. We took the union of the top 2,000 genes with the highest expression and dispersion from both datasets, which were used for PCA. We then aligned the subspaces on the basis of the first 20 PCs and selected top 10 neighbors to report for each batch, which generated new PCs that were used for subsequent analysis.

### Differential expression analysis

To find DEGs among different clusters, we performed non-parametric Wilcoxon rank sum tests, as implemented in Seurat. DEGs with adjusted *P* value less than 0.01 were thought to be significant. We applied tl.rank_genes_groups function with default parameters and DEGs were filtered with “min_fold_change” = 0.5, “min_in_group_fraction” = 0.2, as implemented in Scanpy.

### Surface markers and TFs

Surface markers and TFs lists (**Table S6**) were downloaded from Cell Surface Protein Atlas (http://wlab.ethz.ch/cspa) and HumanTFDB3.0 (http://bioinfo.life.hust.edu.cn/HumanTFDB).

### Correlation analysis

Pearson correlation analysis was performed using the top 50 PCs calculated in CS 13 10X-derived dataset (**Figure S1C**). For genes positively correlated with *RUNX1*, we performed Pearson correlation using corr.test function in psych R package with “BH” adjustment (**Figure 2H**).

### Monocle 2 analysis

Pseudotime trajectories were constructed with the Monocle 2 package (v2.10.1)^86^ according to the documentation (http://cole-trapnell-lab.github.io/monocle-release). ordering genes were selected based on PCA loading. The Discriminative Dimensionality Reduction with Trees (DDRTree) method was used to reduce data to two dimensions. To investigate the different patterns of gene expression during this developmental path, significant DEGs along pseudotime were identified by Monocle 2’s differentialGeneTest function. For identification of major patterns, we filtered out those genes with average normalized expression less than 1 in all involved clusters and then clustered the retained genes into four distinct patterns using k-means clustering.

### Gene functional annotation analysis

GO term and KEGG pathway enrichment was done on significant DEGs using clusterProfiler (version 3.10.1)^87^ with default parameters.

### Cell cycle regression

To reduce the variation in cell cycle stages which contribute to the heterogeneity in scRNA data, we performed CellCycleScoring function in Seurat to evaluate the cell-cycle status using the previously reported G1/S and G2/M phase-specific genes ^81, 88^. Then using regressout in ScaleData function to remove cell cycle effects.

### Cellular interactions analysis

Cellular interactions analysis was performed using CellphoneDB software (version 2.0)^89^ with default parameters. Significant ligand-receptor pairs were filtered by *P* value less than 0.01. Pairs that ligand genes expressed in Mes, Epi, EC clusters and receptor genes expressed in HEC were retained. Directed network (**Figure 5A**) visualization was done using Cytoscape (version 3.6.0).

## DATA AND SOFTWARE AVAILABILITY

The scRNA-seq data has been uploaded to GEO, the accession number for the data is pending.

## DECLARATION OF INTERESTS

The authors declare no competing interests.

## Supplementary Figures and Legends

**Figure S1.**
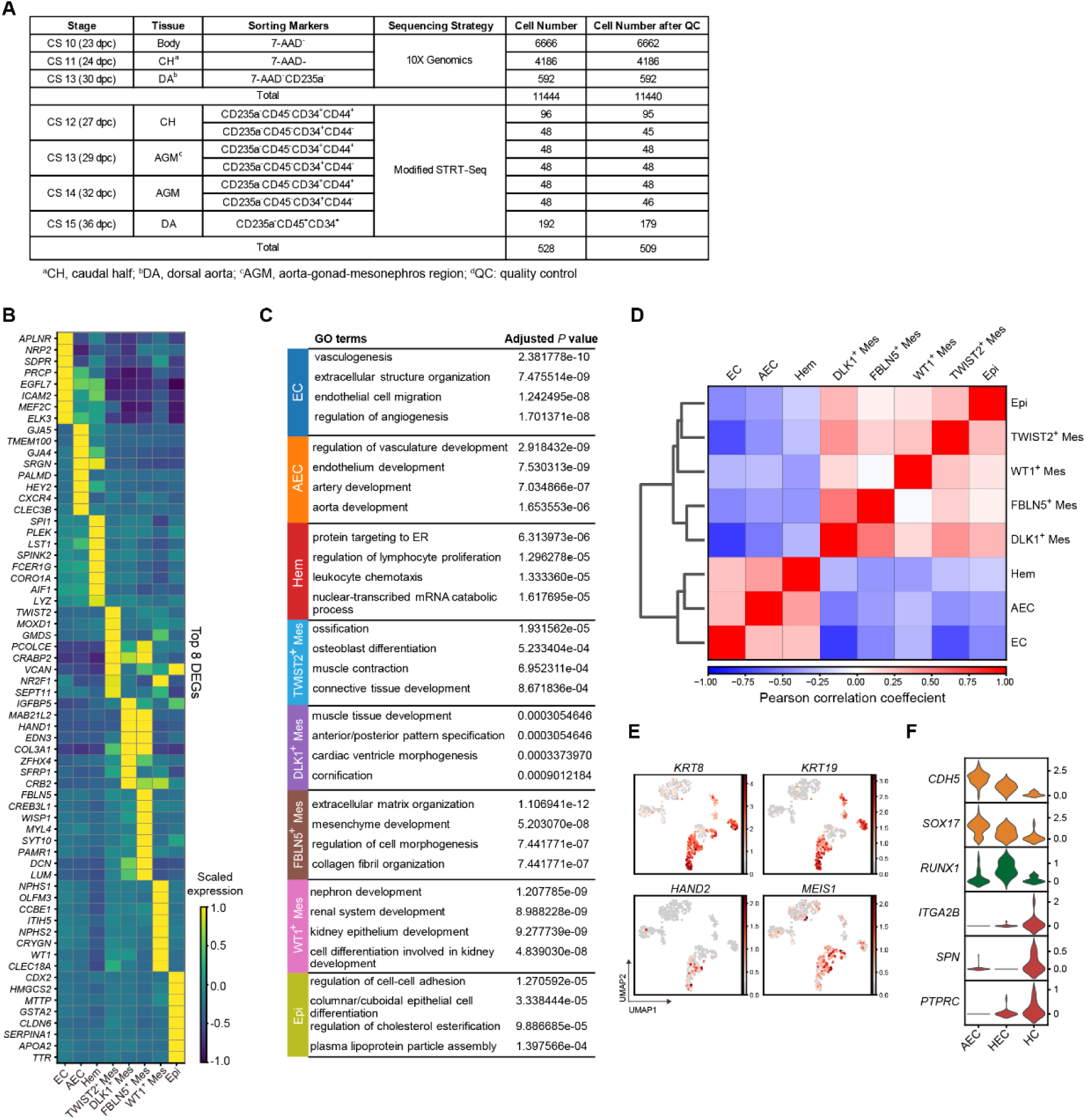
Detailed feature of transcriptomically defined clusters in CS 13 DA. **A**. Sample information for scRNA-seq data. **B**. Heatmap showing Top 8 DEGs for eight clusters. **C**. The enriched major GO:BP terms for each cluster. **D**. Correlation analysis of eight clusters showing that AEC and Hem clusters are more similar with each other, while four mesenchymal populations and Epi cluster correlate more closely to each other. **E**. UMAP plots showing the expression of featured genes of stromal cells. **F**. Violin plots showing the expression of feature genes in AEC, HEC and HC.

**Figure S2.**
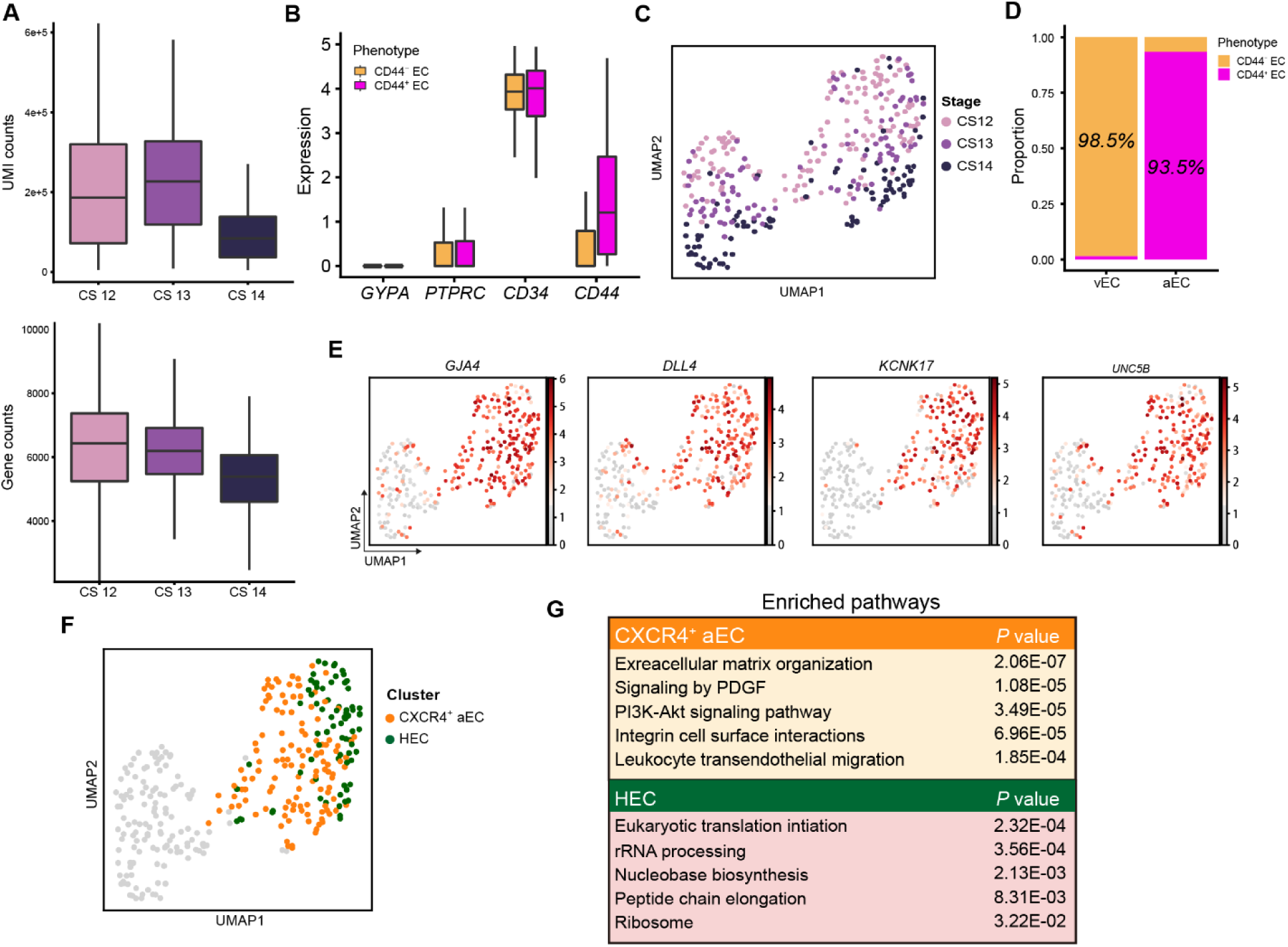
Quality of scRNA-seq data of CD44^+^/CD44^−^ ECs and feature of two sub-divided aEC clusters. **A**. Box plots displaying UMI counts and gene counts of scRNA-seq data generated from CD44^+^/CD44^−^ ECs of CS12 CH, CS13 and CS14 AGM region. Sorting strategy for sequencing is displayed in Figure 2A. **B**. Box plots for the expression of surface markers (CD235a (encoded by *GYPA*) and CD45 (encoded by *PTPRC*)) used for FACS sorting. **C.** UMAP plots for stages of cells showed in Figure 2B. **D**. Bar plots showing the proportion of CD44^+^ EC and CD44^−^ EC in aEC and vEC. **E**. UMAP plots showing the expression of arterial genes in aEC and vEC clusters. **F**. Mapping of CXCR4^+^ aEC and HEC in populations depicted in Figure 2B. **G**. Pathways enriched in two aEC sub-clusters.

**Figure S3.**
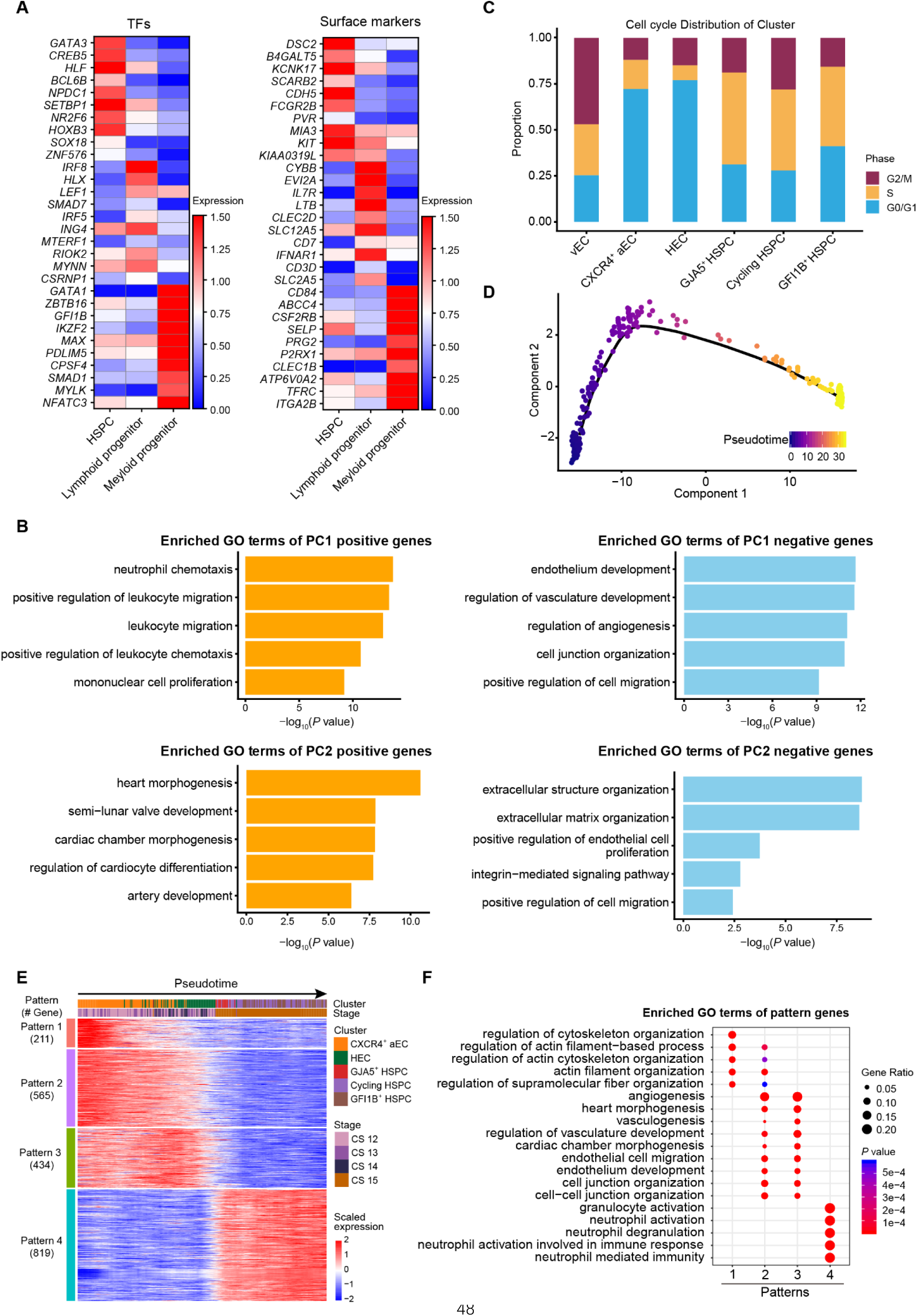
Different patterns of gene expression and cell cycle status during endothelial-to-hematopoietic transition. **A**. Heatmaps of top 10 differentially expressed TFs (left panel) and surface markers (right panel) in HSPC (HSPC1/2/3), Myeloid progenitor and Lymphoid progenitor clusters. **B**. The enriched GO:BP terms for PC 1/2 positive/negative genes are shown, corresponding to the property of vEC, CXCR4^+^ aEC, HEC and three HSPC clusters as shown in Figure 3F. **C**. Cell cycle distribution of vEC, CXCR4^+^ aEC, HEC and three HSPC clusters (GJA5^+^ HSPC, Cycling HSPC and GFI1B^+^ HSPC). **D**. Pseudotime analysis by Monocle 2. **E**. Heatmap showing four different expression patterns along the development of arterial ECs via HSC-primed hemogenic ECs to HSPCs. **F**. The enriched GO terms of four different expression patterns.

**Figure S4.**
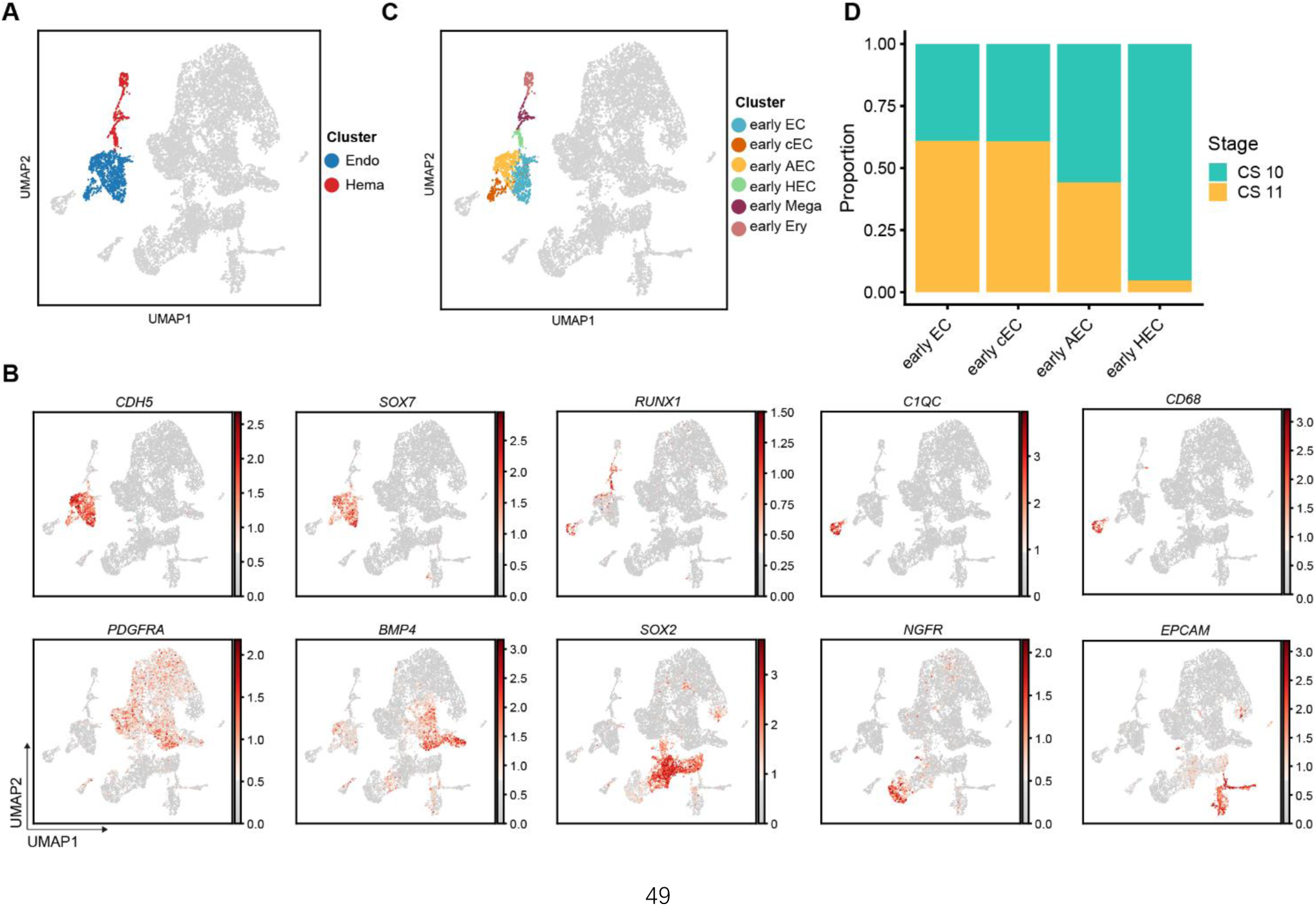
Identification of populations in early stages (CS 10 and CS 11) **A**. Mapping of endothelial (Endo) and hematopoietic cells (Hema) in CS 10 and CS 11. Other populations that do not show a direct relationship to Endo and Hema are shown in gray. **B**. The expression of endothelial, hematopoietic and mesenchymal genes that can help to identify populations in **Figure S4A**. **C**. Mapping of clusters shown on **Figure 4A**. **D**. Bar plot showing ratios that cells at two stages take in early HEC depicted in **Figure 4F**.

**Figure S5.**
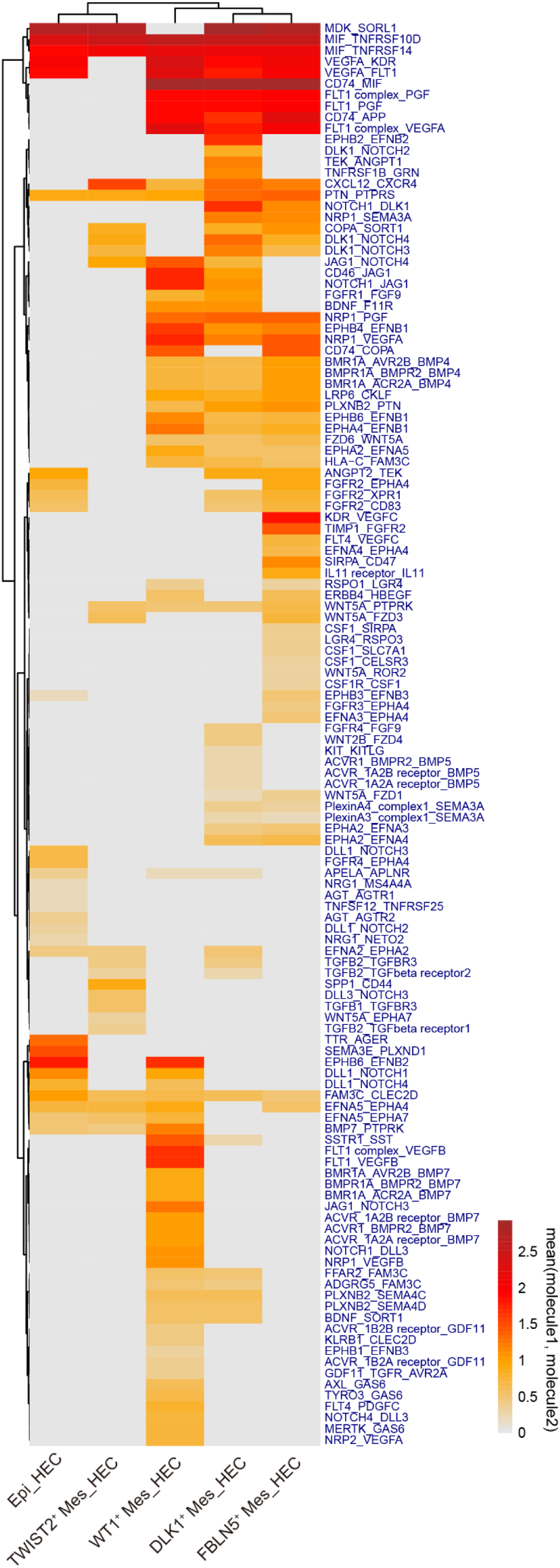
The cell-cell interaction involved in the development of HSC-primed hemogenic ECs predicted by scRNA-seq data. Heatmap showing the detailed ligand-receptor pairs exhibiting different expression patterns among distinct stromal populations when coupled with HEC.

